# Structure of Human BCCIP and Implications for Binding and Modification of Partner Proteins

**DOI:** 10.1101/2020.12.08.416925

**Authors:** Woo Suk Choi, Bochao Liu, Zhiyuan Shen, Wei Yang

## Abstract

BCCIP was isolated based on its interactions with tumor suppressors BRCA2 and p21. Knockdown or knockout of BCCIP causes embryonic lethality in mice. BCCIP deficient cells exhibit impaired cell proliferation and chromosome instability. BCCIP also plays a key role in biogenesis of ribosome 60S subunits. BCCIP is conserved from yeast to humans, but it has no discernible sequence similarity to proteins of known structures. Here we report two crystal structures of an N-terminal truncated human BCCIPβ, consisting of residues 61-314. Structurally BCCIP is similar to GCN5-related acetyltransferases (GNATs) but contains different sequence motifs. Moreover, both acetyl-CoA and substrate-binding grooves are altered in BCCIP. A large 19-residue flap over the putative CoA binding site adopts either an open or closed conformation in BCCIP. The substrate binding groove is significantly reduced in size and is positively charged despite the acidic isoelectric point of BCCIP. BCCIP has potential binding sites for partner proteins and may have enzymatic activity.

## 1 INTRODUCTION

BCCIP is a nuclear protein that was identified in human genome based on its interactions with tumor suppressors BRCA2 and p21^1, 2^. In humans, there are two isoforms resulting from alternative RNA splicing, BCCIPα (322 residues) and BCCIPβ (314 residues), which share the identical N-terminal 258 residues but differ in the remaining C-terminal regions^1, 2^. BCCIPα and β are also known as TOK-1a and TOK-1 β, respectively^1^. BCCIPβ is the conserved isoform in eukaryotes, from yeast, worms, plants to mammals^2^, while BCCIPα only exists in humans. In mouse, there is only one BCCIP, which is ~70% identical to human BCCIPβ. Either knockdown or knockout of BCCIP in mice leads to embryonic lethality due to impaired cell proliferation^3, 4^. BCCIP deficient mouse embryo fibroblast cells exhibit increased sensitivity to DNA damage and replication stress and increased chromosome instability, including chromosome breaks and sister chromatid union^3^. The yeast homolog of BCCIPβ, known as BCP1, appears to be involved in nuclear export and ribosome biogenesis^5, 6^. Recently, both mouse BCCIP and human BCCIPβ were shown to be located in the nucleolus and required for rRNA maturation and ribosome 60S subunit biogenesis^7^, but with distinct features from the yeast BCP1.

Despite its functional importance, structures of BCCIP-family proteins remain elusive. Except for limited sequence similarity of ~50 residues to the Ca^2+^-binding domain in calmodulin and M-calpain^2^, BCCIP has no discernible sequence homology to any known protein. Using isomorphous replacement we have determined crystal structures of a large fragment of human BCCIPβ (aa 61-314) in two different crystal forms. To our surprise, structurally BCCIP is homologous to the GCN5-related N-acetyltransferases (GNAT s)^8^, which use acetyl-CoA to modify primary amines of lysine sidechains or protein N termini, as well as aminoglycosides (antibiotics) and hormones (serotonin)^9–12^. The regions corresponding to the substrate and catalytic motifs in N-acetyltransferases (motifs A-D), are also conserved among BCCIP-family members but are different from GNATs. Whether BCCIP is an enzyme, and what its substrates might be are unclear. While we were engaged in trying to figure out its structure-function relationship, a yeast BCP1 structure was reported earlier this year (with the structure coordinates on hold)^13^. Here we present crystal structures of the conserved isoform of human BCCIPβ and its outstanding binding surfaces for partner proteins and possibly for small molecules.

## 2 RESULTS AND DISCUSSION

### 2.1. Crystal structures of human BCCIP

We have determined crystal structures of N-terminal truncated BCCIPβ (aa 61-314, referred to as BCCIP hereafter) in two different space groups at resolutions of 3.06 (Native1) and 2.13 Å (Native2), respectively (Table 1). The two structures are superimposable except for a 19-residue extended and flexible loop. BCCIP forms a single domain of α/β fold, including a mixed seven-stranded β sheet surrounded on either side by four and three long α helices (Fig. 1). The structure can be divided into two halves. The N-terminal half (aa 61-185) folds into two βααβ hairpins with the four β strands (β1-β4) forming an antiparallel sheet, and four α helices forming two hairpins (αA-αB, and αC-αD) covering one face of the β sheet. Helix αC is preceded by a short helix αC’ (aa132-136). The C-terminal half (aa 186-314) folds in the order of αβαββα and forms a three-stranded antiparallel β sheet (β5-7), with three α helices (αE-αG) on the side opposite to helices αA-αD (Fig. 1a-b). The two halves are linked by helix αE and form a contiguous β sheet by hydrogen bonds at the beginning of parallel strands β4 and β5. These two strands diverge at the end giving the β sheet a V shape (Fig. 1b). No Ca^2+^-binding module or metal ion-binding site can be found. The extended loop linking β6 and β7, L_67_ (aa 269-287), has two dramatically different conformations. In Native1 L_67_ is extended and interacts with a crystallographic symmetry mate reciprocally. But in Native2 L_67_ is folded next to αA-αB, and L_67_ and αAB are like two arms embracing the β sheet (Fig. 1b). The N-terminal half followed by αE and β5 has the same sequence in the two isoforms of human BCCIP, and only αF-β6-β7-αG differ between the two.

**Figure 1.**
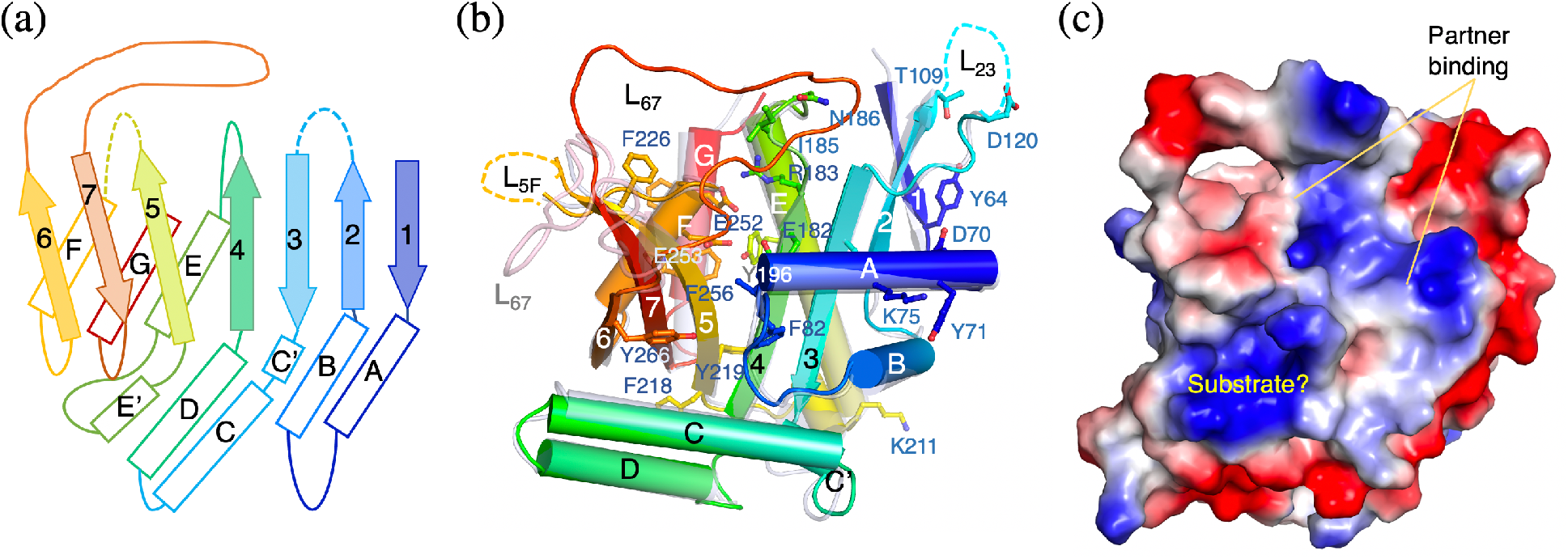
Crystal structures of BCCIP. (a) Topology diagram of BCCIP. The disordered loops are indicated by dashed lines. (b) Cartoon diagrams of superimposed Native1 (semi-transparent grey and pink) and Native2 (solid rainbow colors) structure of BCCIP. Conserved residues are shown as sticks and labeled. (c) Molecular surface of BCCIP (Native2) with electrostatic potential in the same view as in panel b. The potential substrate-binding pocket and interface for protein partners are indicated.

**Table 1.** Statistics of crystallographic data collection and structure refinement

| **Crystal** | Hg derivative | Native1 | Native2 |
| --- | --- | --- | --- |
| PDB |  | 7KYQ | 7KYS |
| Space group | P 4<sub>1</sub>2<sub>1</sub>2 | P 4<sub>1</sub> 2<sub>1</sub> 2 | P 2<sub>1</sub> |
| Unit cell |  |  |  |
| *a*, *b*, *c* (Å) | 114.4, 114.4, 53.9 | 112.7, 112.7, 56.4 | 60.0, 67.0, 99.5 |
| $\alpha$, $\beta$, $\gamma$ (°) | 90.0, 90.0, 90.0 | 90.0, 90.0, 90.0 | 90.0, 91.0, 90.0 |
| Resolution (Å) <sup>a</sup> | 40.0 - 3.62 (3.68 - 3.62) | 40.0 - 3.04 (3.09 - 3.04) | 30.0 - 2.13 (2.19 - 2.13) |
| Wavelength (Å) | 1.0000 | 1.0000 | 1.0000 |
| *R*<sub>merge</sub> (%) <sup>a</sup> | 11.0 (86.7) | 10.7 (98.4) | 9.1 (157.9) |
| *I* / $\sigma$ <sup>a</sup> | 22 (17) | 26 (2.3) | 7.8 (1.0) |
| Completeness (%) <sup>a</sup> | 100.0 (100.0) | 99.8 (99.4) | 98.9 (99.9) |
| Multiplicity <sup>a</sup> | 20.2 (21) | 12.4 (12.1) | 3.7 (3.6) |
| **Refinement** |  |  |  |
| Resolution (Å) <sup>a</sup> |  | 39.8 – 3.06 | 29.7 – 2.13 |
| No. reflections |  | 6877 | 43893 |
| No. reflections for *R*<sub>free</sub> |  | 367 | 1001 |
| *R*<sub>work</sub> / *R*<sub>free</sub> |  | 0.219 / 0.251 | 0.209 / 0.234 |
| No. atoms |  |  |  |
| Overall |  | 1894 | 5767 |
| Protein |  | 1850 | 5490 |
| Ligand |  | - | 39 |
| Water |  | 44 | 238 |
| Mean *B*-factors |  |  |  |
| Overall |  | 86.8 | 69.5 |
| Protein |  | 86.9 | 69.5 |
| Ligand |  | - | 76.3 |
| Water |  | 84.1 | 69.2 |
| R.M.S. deviations |  |  |  |
| Bond lengths (Å) |  | 0.013 | 0.009 |
| Bond angles (°) |  | 1.539 | 1.262 |
| Ramachandran |  |  |  |
| Favored (%) |  | 95.54 | 99.4 |
| Allowed (%) |  | 4.46 | 0.6 |
| Outliers (%) |  | 0 | 0 |

<sup>a</sup> Values in parenthesis are of the highest resolution shell.

### 2.2. BCCIP is structurally similar to GNAT acetyltransferases

Structural similarity search by Dali revealed that BCCIP is similar to GCN5-related acetyltransferases (GNATs), with substrates ranging from histones to antibiotics and hormones. The N-terminal half of BCCIP is superimposable with these GNATs (Fig. 2a). But in GNATs helices αC and αD are absent and β3 and β4 are linked by a β hairpin directly or a random coil^9–12, 14^. Helices αC and αD effectively close the open binding groove for protein substrates, such as the histone tail bound to GCN5^14^, and fill it with hydrophobic residues. With its positively charged pocket (Fig. 1c) BCCIP may still be able to bind small molecules, such as antibiotics and hormones.

**Figure 2.**
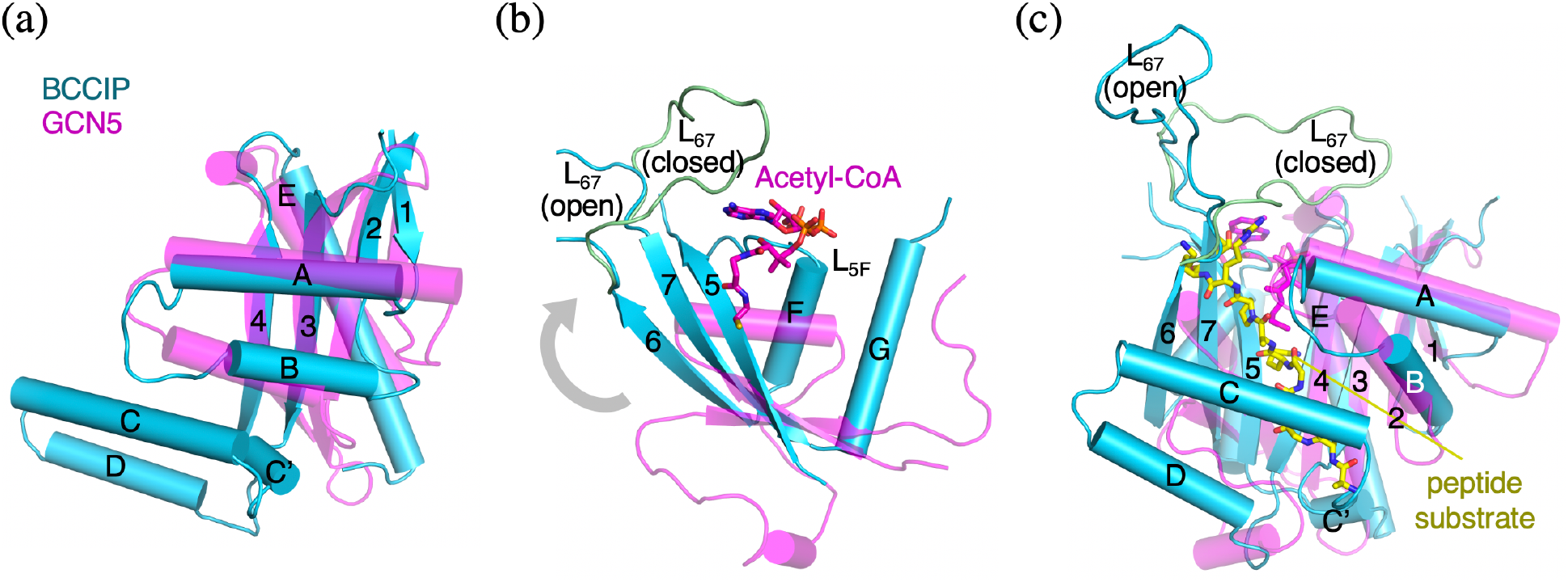
Structural comparison with GCN5. (a) Superposition of the N-terminal half of BCCIP and GCN5 (PDB: 1QSN). (b) After superimposing the N-terminal half, the C-terminal half of BCCIP appears rotated by over 90° (indicated by the grey arrowhead) relative to that of GCN5. Loop L_5F_ in BCCIP would clash with acetyl-CoA (bound to GCN5) if present. (c) The entire BCCIP structure is superimposed with GCN5 via the N-terminal half. Histone peptide bound to GCN5 is shown in a yellow (carbon) stick model.

The structurally most conserved region (motifs A and B) involving in acetyl-CoA binding in GNATs is located between two halves of the structure at β4-β5^8^. The corresponding region is also conserved among BCCIP-family members, but the sequence motifs of BCCIP (Fig. 1b) differ from GNATs^15^.

The C-terminal half of BCCIP is oriented differently from GNATs, particularly loop L_5F_, which binds the diphosphates of CoA in GNATs. BCCIP contains an insertion in L_5F_, and without structural changes, its L_5F_ would clash with the CoA moiety Fig. 2b). In case substrate binding induces conformational changes in BCCIP, we tried co-crystallization of BCCIP with 10 different acyl-CoA compounds. Instead of finding a bound substrate, we obtained a different form of apo BCCIP crystal (Native2) (see section 3.2).

In addition, compared to GNATs, BCCIP has the following two insertions, the extended flexible loop L_67_, which is open and extended in Native1 and adopts a closed conformation in Native2 (Fig. 2c), and the C-terminal helix αG. Together with L_5F_, these insertions alter the acetyl-CoA binding site significantly. Peculiarly, two Tyr residues, Y64 and Y71, are exposed

### 2.3. Potential binding surface for protein and small molecules

The conserved residues among BCCIP homologs are clustered into two groups. One is around β1-αA and the loop linking β2 and β3, forming a predominantly negatively charged convex surface (Fig. 1c). Peculiarly two conserved Tyr, Y64 and Y71, are solvent exposed and begging for an interacting partner. The other is between β4 and β5, the V-shaped substrate-binding groove and pocket (top and front in Fig. 1c). Aromatic residues also abound the second cluster. Surrounding the V-shaped binding groove, two disordered loops L_23_ (aa 110-119) and L_5F_ (aa 229-244) are next to the flexible flap L_67_ (Fig. 1a-b), like tentacles waiting to embrace a catch. Human, mouse and yeast BCCIPs are overall negatively charged with isoelectric point below 4.8. However, substrate binding groove and pocket have a positively charged appearance, but they are surrounded by an intensely negatively charged belt (Fig. 1c). These features indicate that BCCIP binds specific protein partners and possibly a small molecule substrate in the same location as GNATs.

## 3 MATERIALS AND METHODS

### 3.1. BCCIP expression and purification

The N-terminal 60 residues of BCCIPβ were predicted to be highly disordered, and probably prevented crystallization of the full-length BCCIPβ (data not shown). The N-terminal truncation construct of BCCIPβ (aa 61-314, abbreviated as BCCIP) was cloned into pET-28c(+) (Novagene Corp. Inc.) between NcoI and EcoRI sites, and the resulting plasmid was transformed into BL21 (DE3) competent cells. The cells were grown in LB medium at 37°C until OD_600_ reached ~0.6. After cooling down to 16°C for 30 min, expression of BCCIP was induced by addition of 0.3 mM IPTG at 16 °C for 20 hours. The cells were pelleted, re-suspended in buffer A (20 mM sodium phosphate (pH7.4), 500 mM NaCl) with 40 mM imidazole, and lysed by sonication. The lysate was cleared by centrifugation at 30,000g for 1 hour at 4°C. BCCIP was bound to an Ni^2+^ column and eluted in buffer A plus 300 mM imidazole. The eluted protein was further purified using a Hitrap Q HP column (GE Healthcare) equilibrated in buffer B (25mM Tris-HCl (pH 7.5), 100 mM NaCl, 1 mM EDTA, 0.01% IGEPAL-CA630, and 1 mM DTT) and was eluted with a NaCl gradient from 100 to 1000 mM. As a final step, BCCIP was applied to a Hiprep Sephacryl S-200 16/60 column (GE healthcare) equilibrated in buffer B. The BCCIP peak was pooled and concentrated to ~15 mg/ml and stored at −80 °C in buffer B and 50% glycerol.

### 3.2. Crystallization

BCCIP was buffer exchanged to 25 mM Tris-HCl (pH 7.5) and 1 mM DTT in an Amicon Ultra-0.5 (Millipore, 10k cutoff) and concentrated to 15 mg/ml (514 μM). Crystallization screening was performed at 4°C with JCSG core I (Qiagen) using the sitting drop vapor diffusion method. The protein and precipitant were mixed at equal volume (0.1 *μ*l each). Native1 crystals were obtained in 0.2 M tri-sodium citrate and 18-20% PEG3350. The initial crystals diffracted X-rays only to ~3.3 Å. After improving protein purification and additions of 1 mM DTT, 100 mM NaCl and 50% glycerol in the protein storage buffer, BCCIP crystals (Native1) became much more reproducible and diffracted X-rays to nearly 3.0 Å.

After realizing that BCCIP is similar to GNATs, we tested co-crystallization of BCCIP with various acyl-CoA compounds^16^. Among 10 acyl-CoAs we tried, crystals grown with 1.1 mM benzoyl(bz)-CoA with a precipitant of 0.1 M citric acid (pH 4.5) and 1-2% PEG 6000 at 20°C diffracted X-rays to 2.13 Å. But after structural determination by molecular replacement, the resulting electron density map showed no bz-CoA. Bz-CoA thus functioned as an additive in crystallization, and it changed the crystal space group with improved X-ray diffraction, to a form we termed Native2 (Table 1).

The BCCIP crystals were transferred into the respective stabilization solutions (Native1 in 0.2 M tri-sodium citrate (pH 8.0), 25% PEG 3350, and 1 mM DTT, Native2 in 0.1 M citric acid (pH 4.5), 5% PEG 6000, and 1 mM DTT) for 30 min before freezing. Native2 crystals cracked into smaller pieces in the stabilization solution. BCCIP crystals were cryo-protected by 20% (Native1) or 3 % (Native2) ethylene glycol and flash-frozen in liquid nitrogen.

### 3.3. Phase determination

We selected twelve commonly used Hg (mercury), Pt (platinum), Au (gold) and Pb (lead) compounds, each of which was tested in 0.2 mM or 5 mM concentration, and soaked into the crystals for 1 to 24 hours. We monitored crystal morphology, birefringence and X-ray diffraction at various time intervals. Before flash-freezing in liquid nitrogen, crystals were washed in the stabilization buffer for 15-20 second. R_iso_ between a heavy-atom soaked and native crystal was calculated using Scaleit implemented in CCP4i^17, 18^ to evaluate whether a derivative was made. We determined the BCCIP structure with phases obtained from six datasets of ethylmercuri-thiosalicylic acid derivative (3.15-4.2 Å, Table 1).

### 3.4. Data collection and structure determination

All X-ray data were collected at beamline 22ID (Native1) and 22BM (Native2 and Hg-derivative) at the Advanced Photon Source (APS) at Argonne National Laboratory. Each dataset was processed using HKL2000 (Native1 and Hg derivative)^19^ and XDS (Native2)^20^. Native1 crystals were in P4_1_2_1_2 space group with one molecule per asymmetric unit, and Native2 crystals were in P21 space group with three molecules per asymmetric unit (Table 1). For SAD (Single-wavelength Anomalous Dispersion) phasing, 6 Hg-derivative datasets were merged to enhance the anomalous signal using HKL2000 at 3.6 Å resolution (Table 1)^19^, and phases were calculated using AutoSol in Phenix^21^. The electron density map was further improved by phase extension to 3.04 Å with the native data. An initial model was built using AutoBuild in PHENIX^21^. Because diffraction of Native1 crystals was highly anisotropic, we corrected the data scaling and truncation using Diffraction Anisotropy Server^22^ to improve the map and refinement. Aided by the secondary structure prediction by PsiPred^23^ we manually traced 228 out of 254 residues using COOT^24^ and refined the BCCIP structure using PHENIX^21^. (Table 1). Two loops linking β2 to β3 (L_23_, aa 110-119) and β5 to αF (L_5F_, aa 229-244) were disordered (Fig. 1).

Native2 structure was determined by molecular replacement using Phaser in PHENIX^21^ with Native1 BCCIP structure as the search model. Model building and refinement were carried out as described above, and non-crystallographic symmetry constraints were applied. Except for the re-orientation of the long loop L_67_ (aa 269-287) due to different crystal lattice contacts, the two BCCIP structures were superimposable (Fig. 1).

All structure figures were prepared with PYMOL (www.pymol.org).

## ACKNOWLEDGEMENTS

The authors thank Dr. M. Gellert for critical reading of the manuscript. The research was supported by the NIH intramural research funding to W. Y. (DK036146) and NIH grant (R01CA195612) to Z.S.

## AUTHOR CONTRIBUTIONS

**Bochao Liu**: BCCIP preparation and initial crystallization screen. **Woo Suk Choi**: Crystallization, structure determination, analysis and validation, and manuscript preparation. **Wei Yang**: structure analysis, figure preparation and writing the original draft. **Zhiyuan Shen**: Conceptualization; investigation, writing and editing.

